# Epistatic impacts of *cis*- and *trans*-regulatory mutations on the distribution of mutational effects for gene expression in *Saccharomyces cerevisiae*

**DOI:** 10.1101/2025.03.26.645465

**Authors:** Eden McQueen, Bing Yang, Patricia J. Wittkopp

## Abstract

Epistasis can influence evolution by causing the distribution of phenotypic effects for new mutations to vary among genotypes. Here, we investigate how epistatic interactions between new mutations and an existing regulatory mutation might impact the evolution of gene expression using *Saccharomyces cerevisiae.* We do so by estimating the distribution of mutational effects for expression of a fluorescent reporter protein driven by the *S. cerevisiae TDH3* promoter in a reference strain as well as in eight mutant strains. Each of the mutant strains differed from the reference strain by a single mutation affecting expression of the focal gene. We found that half of these regulatory mutations changed the variance and/or skewness of the distribution of mutational effects. A change in variance indicates a change in mutational robustness, and we found that one initial regulatory mutation increased mutational robustness while another decreased it. A change in skewness indicates a change in the relative frequency and/or effect size of mutations increasing or decreasing expression, and we found that the initial regulatory mutation in four strains had such an effect. Strikingly, in all four of these cases, the change in skewness increased the likelihood that new mutations would at least partially compensate for the effects of the initial regulatory mutation. If this form of epistatic impact on the distribution of mutational effects is common, it could provide a neutral mechanism reducing the divergence of gene expression and help explain the prevalence of alleles with compensatory effects in natural populations of *S. cerevisiae*.

## Introduction

Mutations provide the raw material for evolution by generating the phenotypic variation upon which natural selection can act. For any given genotype, the effects of all possible new mutations on a focal phenotype are referred to as the distribution of mutational effects for that phenotype. This distribution is important because it defines the available phenotypic space for a subsequent evolutionary step. Factors that change the shape of the distribution of mutational effects can therefore influence evolution by making it more or less likely that new mutations produce specific phenotypes (Eyre-Walker and Keightley 2007; Silander et al. 2007). Knowing what these factors are and the nature of their influence on the distribution of mutational effects is informative when making evolutionary predictions.

Epistasis (non-additive interaction among genetic variants at different loci) is one factor that is known to influence the phenotypic effects of new mutations (Phillips 2008; Hansen 2013; Johnson et al. 2023). Indeed, theoretical work (Phillips 2008; Hansen 2013) and empirical studies (e.g., (Bridgham et al. 2006; Weinreich et al. 2006; Salverda et al. 2011; Lang and Desai 2014) have shown that epistasis is capable of impacting the trajectory of phenotypic evolution. Studies focusing on fitness have shown that epistasis can impact the distribution of mutational effects for fitness (also known as the distribution of fitness effects) by altering the distribution’s skewness and/or variance (Silander et al. 2007; Kryazhimskiy et al. 2014; Wünsche et al. 2017; Aggeli et al. 2021; Johnson and Desai 2022; Ascensao et al. 2023; Couce et al. 2024). Interestingly, such studies often find evidence of “diminishing returns” epistasis, wherein the frequency and/or effect size of beneficial mutations decreases as the fitness of the starting genotype increases (Reviewed in Johnson et al. 2023). This observation suggests that–at least in the case of *fitness*–epistasis can have some predictable influences on evolutionary trajectories despite the idiosyncratic epistatic effects observed between particular pairs or sets of mutations (Kryazhimskiy et al. 2014; Reddy and Desai 2021; Bakerlee et al. 2022); (Johnson et al. 2023).

But what about traits other than fitness? We currently know very little about how epistatic interactions among mutations distributed throughout the genome influence the distribution of mutational effects for other types of polygenic traits, including gene expression. Gene expression is regulated by networks of *cis*-acting DNA sequences containing binding sites for *trans*-regulatory proteins and RNAs, many of which themselves interact. Because of this interconnectedness, theoretical studies predict that epistasis should be common between mutations affecting components of gene regulatory networks (Omholt et al. 2000; Gjuvsland et al. 2007; Uller et al. 2018; Fagny and Austerlitz 2021), making gene expression an attractive phenotype for tackling this question.

Thus far, empirical studies testing for epistatic interactions among new mutations affecting a gene’s expression have focused on mutations targeted to a subset of that gene’s regulatory network (e.g., Li et al. 2019; Lagator et al. 2017). For example, Lagator et al (2017) examined the impacts of epistasis on the distribution of mutational effects for gene expression using mutations targeted to a single transcription factor and its binding site in the promoter of a viral gene tested in bacteria. They found that when both functional elements were mutated simultaneously, the phenotypic variance for expression of the focal gene was greater than could be explained by additive interactions between the *cis*- and *trans*-regulatory mutations alone, indicating that epistatic interactions can alter the shape of the distribution of mutational effects for gene expression. In eukaryotes, *cis*-regulatory sequences typically contain multiple binding sites for multiple *trans*-acting transcription factors, and the activity of these transcription factors may be affected by many genes and gene products in the cell, so the epistatic impacts of new regulatory mutations might be complex. Understanding how epistatic interactions alter the effect of new mutations arising throughout the genome is an important empirical target because it provides information about selectively-neutral expectations without making assumptions about the loci that might be involved.

To capture this genome-wide picture of the impacts of epistasis on the distribution of mutational effects for a gene expression phenotype, we measured the effects of new point mutations arising randomly throughout the genome on expression of a fluorescent reporter gene driven by the *S. cerevisiae TDH3* promoter in nine strains of *S. cerevisiae*. Eight of these nine strains were derived from a shared reference strain (i.e., the ninth strain) by engineering a distinct regulatory mutation into each strain. The introduced mutations were previously shown to affect expression of the reporter gene into each strain (Duveau et al. 2014; Metzger et al. 2015; Duveau et al. 2018; Duveau et al. 2021). Four of these regulatory mutant strains contained a single nucleotide change in the focal gene’s promoter (i.e., a *cis*-regulatory element). The remaining four strains contained a non-synonymous mutation in the coding sequence of a gene known to regulate expression of the focal gene in *trans*. The chemical mutagen ethyl methanesulfonate (EMS) was then used to introduce new point mutations into each of these regulatory mutant strains (as well as the reference strain) randomly throughout the genome. Expression of the focal gene was measured in EMS mutants isolated from each strain, and these measurements were used to estimate the strain-specific distribution of mutational effects for our expression phenotype. Differences in the shape of the distribution of mutational effects observed between the reference strain and one of the regulatory mutant strains were attributed to epistatic interactions between the specific regulatory mutation present in that strain and the new mutations introduced by EMS.

We found that one of the *cis*-regulatory mutants and three of the *trans*-regulatory mutants showed a significant change in the variance and/or skewness of the distribution of mutational effects for the focal gene’s expression. One of the *trans-*regulatory mutant strains showed a larger variance than the reference strain, indicating that the regulatory mutation initially engineered into this strain decreased the mutational robustness of the focal gene’s expression relative to the reference strain. A second mutant strain carried a *trans*-regulatory mutation that decreased the variance of the distribution of mutational effects, indicating that it increased mutational robustness. Significant differences in skewness were also observed for the distribution of mutational effects for these two trans-regulatory mutant strains as well as one additional *trans*-regulatory mutant strain and one of the cis-regulatory mutant strains. Strikingly, in all four of these cases, the change in skewness was such that mutations counteracting (i.e. at least partially compensating for) the effects of the initial regulatory mutation had larger average effects than mutations exacerbating the effects of the initial mutation. This type of epistatic impact on the distribution of mutational effects could slow the rate of expression divergence, even in the absence of natural selection.

Taken together, these data show that epistatic interactions between new mutations and as little as one nucleotide change can alter the distribution of effects on gene expression in ways that might impact phenotypic evolution. Although it is not yet clear how often the specific types of epistatic impacts on a distribution of mutational effects we observed for our focal gene will also be seen for other genes, other starting genetic variants, or other species, our work suggests that allowing distributions of mutational effects to vary among genotypes and over evolutionary time is important for building realistic models of regulatory evolution.

## Results and Discussion

### Study system used for quantifying distributions of mutational effects on gene expression

To empirically measure and compare the effects of new mutations on expression of a focal gene, we used the *P_TDH3_-YFP* reporter gene, which contains the *S. cerevisiae TDH3* promoter allele from the commonly used BY lab strain driving expression of a yellow fluorescent protein (YFP), integrated into the *S. cerevisiae* genome at the *HO* locus (Metzger et al. 2016). This *TDH3* promoter comes from the highly expressed *TDH3* gene encoding a glyceraldehyde-3-phosphate dehydrogenase (GAPDH) protein that functions in yeast metabolism and also affects other traits (McAlister and Holland 1985; Ringel et al. 2013; Branco et al. 2014). Previous studies have used this fluorescent reporter gene, coupled with EMS mutagenesis and flow cytometry, to estimate the distribution of mutational effects for *P_TDH3_-YFP* expression in a reference strain (Gruber et al. 2012; Metzger et al. 2016). Specific *cis*- and *trans*-regulatory mutations that caused a statistically significant change in the activity of the *TDH3* promoter driving YFP expression have been identified and engineered individually into the starting reference strain, creating a set of regulatory mutant strains (Duveau et al. 2014; Metzger et al. 2015; Duveau et al. 2018; Duveau et al. 2021). We selected eight of these strains for the current work, including four *cis*-regulatory mutants and four *trans*-regulatory mutants.

### Properties of the cis- and trans-regulatory mutant strains examined

The four *cis*-regulatory mutant strains (**Figure 1**) each differ from the reference strain by only a single point mutation in the *TDH3* promoter sequence of the reporter gene. Mutations in the *tata1* and *tata2* strains are located in the TATA box, 137 bp and 136 bp upstream of the start codon, respectively, and have been reported to decrease expression to about 25% and 60% of the reference strain (Duveau et al. 2018). The mutation in strain *gcr1_bs^m76^*is located 482 bp upstream of the *TDH3* start codon and alters a binding site for the GCR1p transcription factor (Huie et al. 1992). This mutation has been reported to decrease expression to about 20% of the reference strain ((Duveau et al. 2018). Finally, the mutation in strain *rap1_bs^m66^* is located 505 bp upstream of the *TDH3* start codon and alters a binding site for the RAP1p transcription factor (Yagi et al. 1994). It has been reported to decrease expression of the reporter gene to about 60% of the level seen in the reference strain (Duveau et al. 2018).

**Figure 1.**
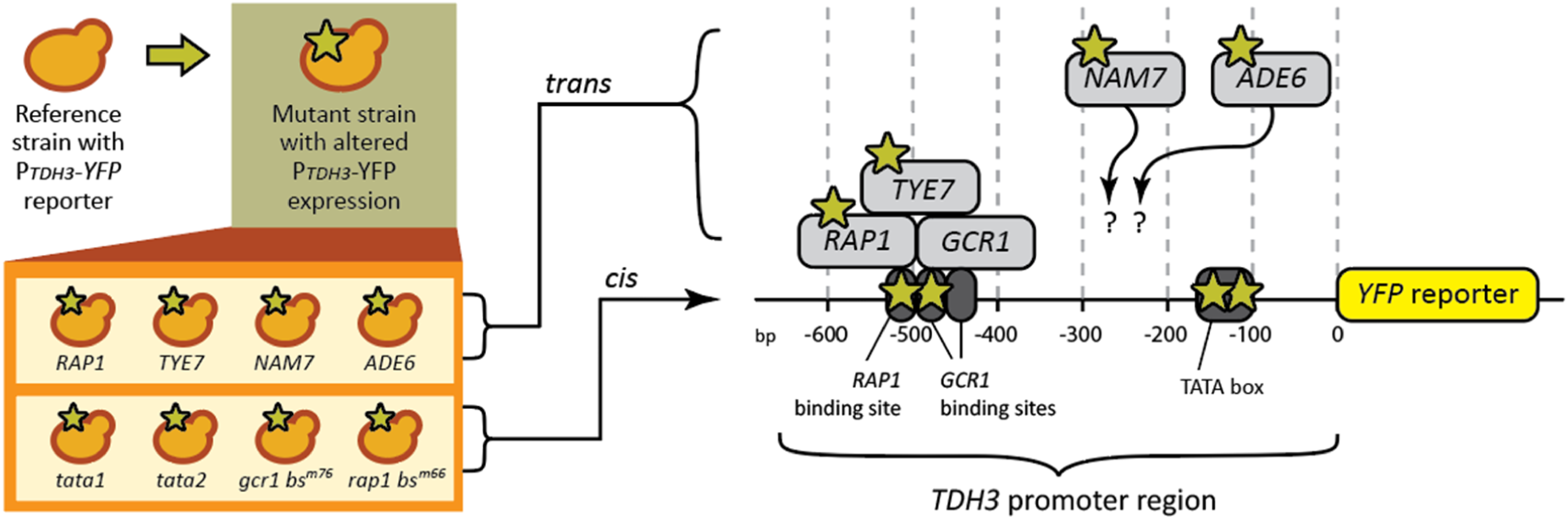
Strains containing *cis*- and *trans*-regulatory mutations affecting expression of the focal gene. The schematic shows a budding yeast cell in the upper left corner that represents the reference strain of *S. cerevisiae* used in this work. This strain contains the *P_TDH3_-YFP* reporter gene integrated into the *HO* locus expressing a yellow fluorescent protein. A generic mutant strain, derived from this reference strain, carrying a mutation that alters YFP expression (yellow star) is also shown. The box connected to this generic mutant contains representations of the eight specific regulatory mutant strains used in this study. The *TYE7*, *RAP1*, *ADE6*, and *NAM7* regulatory mutant strains each contain a non-synonymous point mutation affecting expression of *P_TDH3_-YFP* in *trans* that is located in the coding sequence of the gene the strain is named for. The *tata1, tata2*, *gcr1_bs^m76^*, and *rap1_bs^m66^* strains each contain a point mutation in a functional element (e.g., transcription factor binding site) of the *TDH3* promoter that affects expression of *P_TDH3_-YFP* in *cis.* The image on the right side of the figure shows a schematic of the 678 bp *TDH3* promoter (black line) driving expression of the YFP coding sequence (yellow rectangle) with yellow stars indicating the approximate locations of the mutation present in the *tata1* (-137 bp), *tata2* (-136 bp), *gcr1_bs^m76^*(-482 bp), and *rap1_bs^m66^* (-505 bp) *cis*-regulatory mutant strains. Grey rectangles show proteins with *trans*-regulatory effects on *P_TDH3_-YFP* expression, with yellow stars indicating proteins altered in mutant strains tested. GCR1p and RAP1p are shown binding directly to the *TDH3* promoter, while TYE7p is shown complexing with RAP1p and GCR1p without binding directly to the *TDH3* promoter. ADE6p and NAM7p regulate the *TDH3* promoter indirectly through unknown mechanisms (indicated by question marks).

The four *trans*-regulatory mutants selected each differ from the reference strain by a nonsynonymous mutation in the coding sequence of a gene (*RAP1*, *TYE7*, *NAM7*, or *ADE6*) previously shown to affect *TDH3* expression either directly or indirectly (Duveau et al. 2021) (**Figure 1**). The *RAP1* mutant strain also contains two synonymous mutations in the *RAP1* coding sequence, which are not expected to affect expression of the *P_TDH3_-YFP* reporter gene (Duveau et al. 2021). *RAP1* encodes a transcription factor that directly regulates *TDH3* by binding to a well-established sequence in the *TDH3* promoter (Yagi et al. 1994). *TYE7* also encodes a transcription factor, but regulates *TDH3* indirectly by forming a complex with the RAP1p and GCR1p transcription factors that bind directly to the *TDH3* promoter (Shively et al. 2019)). *NAM7* and *ADE6* encode, respectively, an RNA helicase involved in nonsense-mediated decay (Sheth and Parker 2006) and an enzyme involved in purine biosynthesis (Andreichuk et al. 1997), both of which are assumed to regulate *TDH3* promoter activity indirectly through an unknown mechanism. The nonsynonymous mutations in *RAP1*, *TYE7*, *NAM7*, and *ADE6* cause the unmutated reference *TDH3* promoter allele to drive YFP expression at ∼92%, 80%, 110%, and ∼155% of the level seen in the reference strain, respectively (Duveau et al. 2021).

### Distributions of mutational effects for P_TDH3_-YFP expression in cis- and trans-regulatory mutants

To estimate the distribution of mutational effects for activity of the *TDH3* promoter driving expression of YFP, we first grew a population of cells from each of the eight regulatory mutant strains as well as two replicate populations of cells from the unmutated reference strain (*ref.1* and *ref.2*). For each of these ten populations, half of the cells were treated with the chemical mutagen EMS. The other half of the cells were handled identically except that no EMS was added, which was a control treatment we refer to as a “sham” (**Figure 2A**). EMS introduces mutations randomly throughout the genome, and these mutations are almost exclusively G:C -> A:T transitions (Shiwa et al. 2012; Duveau et al. 2021). These two types of mutations are the most common types of spontaneous point mutations in *S. cerevisiae (Lynch et al. 2008)*. Insertions, deletions, and aneuploidies can also be introduced by EMS or (more likely) arise spontaneously during the experiment, but findings from (Duveau et al. 2021) suggest that these types of mutations should be less than 3% of the mutations sampled. Using a canavanine resistance assay (Lang and Murray 2008; Gruber et al. 2012), we estimated that our EMS treatment introduced an average of 38 new mutations per cell (**Supplementary Table S1**). However, recent genome sequencing of EMS mutants produced using the same mutagenesis protocol found that this canavanine resistance assay overestimated the true number of point mutations by 25% (Duveau et al. 2021), suggesting that each EMS-treated cell isolated in the current study actually carries about 28 mutations. Genetic analysis of the EMS mutants sequenced in the prior study showed that significant changes in *P_TDH3_-YFP* expression were nearly always caused by only one of the point mutations present in an EMS mutant (Duveau et al. 2021), and we expect the same to be true for EMS-treated cells isolated in the current study.

**Figure 2.**
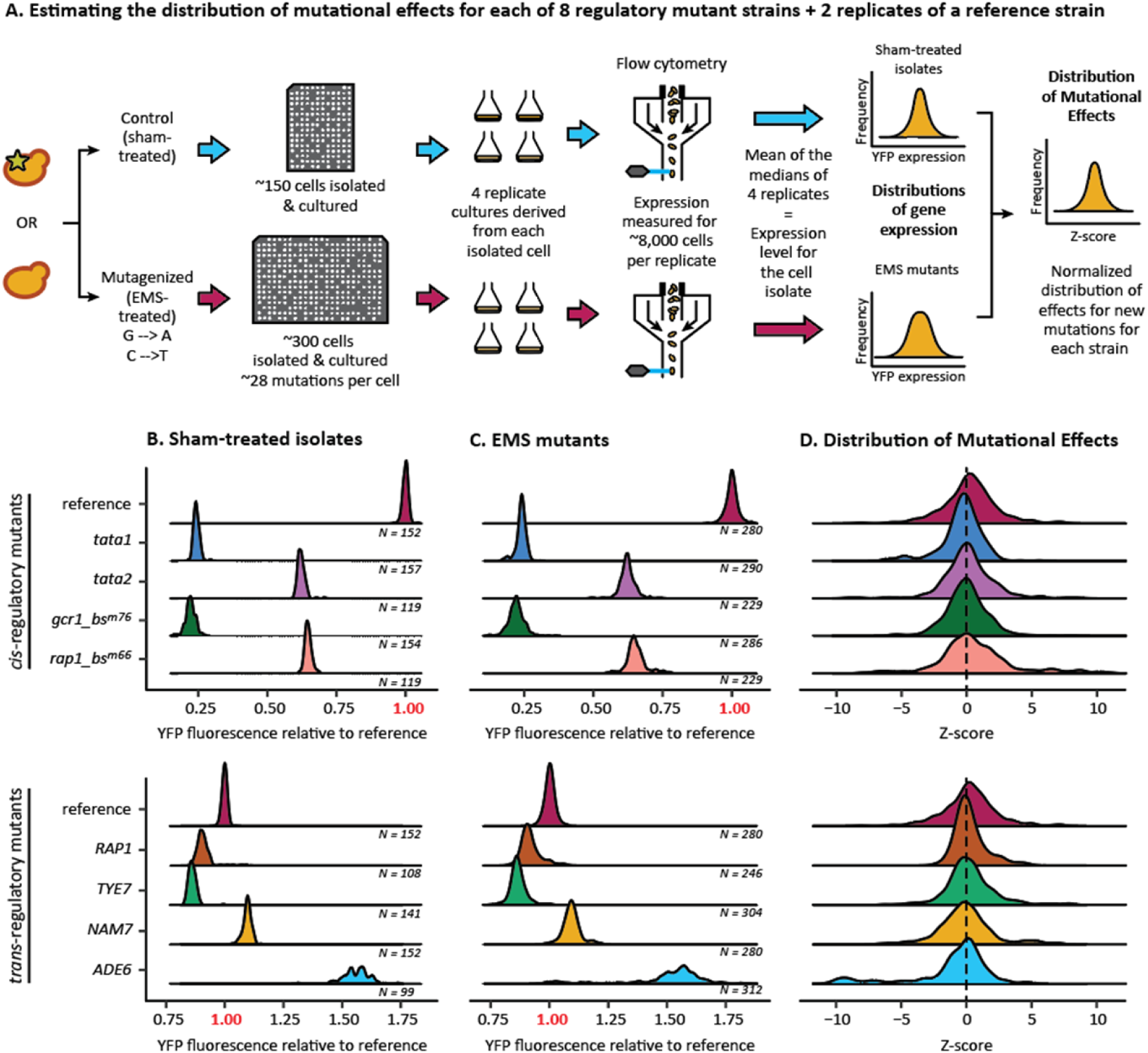
Strain-specific distributions of mutational effects for expression of the focal gene. (**A**) The schematic provides an overview of how we inferred the distribution of mutational effects for each of the eight regulatory mutant strains and two replicates of the reference strain. Each experiment began with a clonal population of genetically identical cells. Some of these cells were exposed to EMS, which introduces G to A and C to T point mutations, and some of the cells were subject to a sham treatment involving all the same steps except for the omission of EMS. Using fluorescence activated cell sorting (FACS), 384 events (cells or similarly sized debris) were randomly sorted from each of the ten EMS-treated populations and deposited individually onto solid plates in a 384-well plate format. The same procedure was followed for each of the ten sham-treated populations, but only 192 events (half of 384) were sorted in each case. After incubating these plates, 229-312 colonies (median = 280) grew from individually sorted EMS-treated cells and 99-157 (median = 147) grew from individually sorted sham control cells (**Supplementary Table S2**). Each colony was used to inoculate four replicate liquid populations, and then fluorescence from the YFP reporter gene was measured in ∼8,000 cells per replicate using flow cytometry. After correcting for variation in cell size, the YFP expression level for each EMS mutant and each sham-treated isolate was estimated by determining the median expression level of cells in each of the four replicate liquid cultures and calculating the mean of those four medians. The mean and standard deviation of expression levels observed among the sham-treated isolates were then used to normalize expression of each EMS mutant isolated from the same starting population by calculating a Z-score. **(B,C)** Distributions of gene expression level (measured as YFP fluorescence relative to the reference strain) observed for the sham-treated isolates (**B**) and EMS mutants (**C**) from each of the four *cis*-regulatory mutants, as well as the *ref.2* sample of the reference strain, are shown in the top half of panels **B** and **C**. Distributions of gene expression level observed for the sham-treated isolates (**B**) and EMS mutants (**C**) from each of the four *trans*-regulatory mutants, as well as the *ref.2* sample of the reference strain, are shown in the bottom half of panels **B** and **C.** Note that the range of YFP fluorescence values shown on the X-axis differs for the *cis*- and *trans*-regulatory mutants to better visualize their effects. The number of EMS mutants and sham-treated isolates analyzed from each strain is shown below each plot. (**D**) Distributions of mutational effects show Z-scores calculated by standardizing YFP expression level in each EMS mutant with the mean and standard deviation of the corresponding sham-treated isolates.

Using fluorescence-activated cell sorting (FACS), 384 randomly chosen “events” (cells or similarly sized debris) from each of the ten mutagenized populations were deposited on a solid plate in a 384-well array (**Figure 2A**). Incubating these plates at 30 °C for 48 hours resulted in growth of 229-312 (median = 280) single colonies from each of the ten mutagenized populations (**Supplementary Table S2**); each colony was assumed to be a distinct EMS mutant. From each sham population, 192 events were similarly sorted and incubated, resulting in 99-157 (median = 147) single colonies for each of the ten starting populations (**Supplementary Table S2**), which were considered sham-treated isolates. Each of the colonies isolated from an EMS- or sham-treated population was used to inoculate four replicate liquid cultures (**Figure 2A**). Using flow cytometry, we then measured fluorescence and cell size for ∼8000 cells from each of these liquid cultures (**Figure 2A**). The *P_TDH3_-YFP* expression level for each EMS mutant and sham isolate was estimated by calculating the mean of the median expression levels (corrected for variation in cell size) from each of the four corresponding replicate liquid cultures (**Figure 2A**).

The distributions of expression levels for the EMS mutants and sham-treated isolates from each of the *cis*- and *trans*-regulatory mutant strains are shown in **Figure 2B** and **Figure 2C**. Distributions of expression levels for *ref.1* and *ref.2* were very similar to each other (**Table 1**). *Ref.2* (chosen by coin flip) is shown in **Figure 2** to represent the reference strain. The mean expression level observed among the sham-treated isolates from each of these strains (**Table 1**, **Figure 2B**) represents that strain’s starting expression level. In all cases, the difference in mean expression level between the sham-treated isolates from a regulatory mutant strain and the *ref.2* sample of the reference strain was similar to the previously described effect of the initial *cis*- or *trans*-regulatory mutation present in that strain (**Table 1**). All sham-treated isolates from the same strain were assumed to be genetically identical because prior work suggested that mutations arising spontaneously during the sham treatment were negligible (Gruber et al. 2012; Metzger et al. 2016). Consequently, the variation seen in each of the sham distributions is a reflection of the expression noise for that strain ((Hornung et al. 2012; Metzger et al. 2015; Duveau et al. 2018).

**Table 1.**
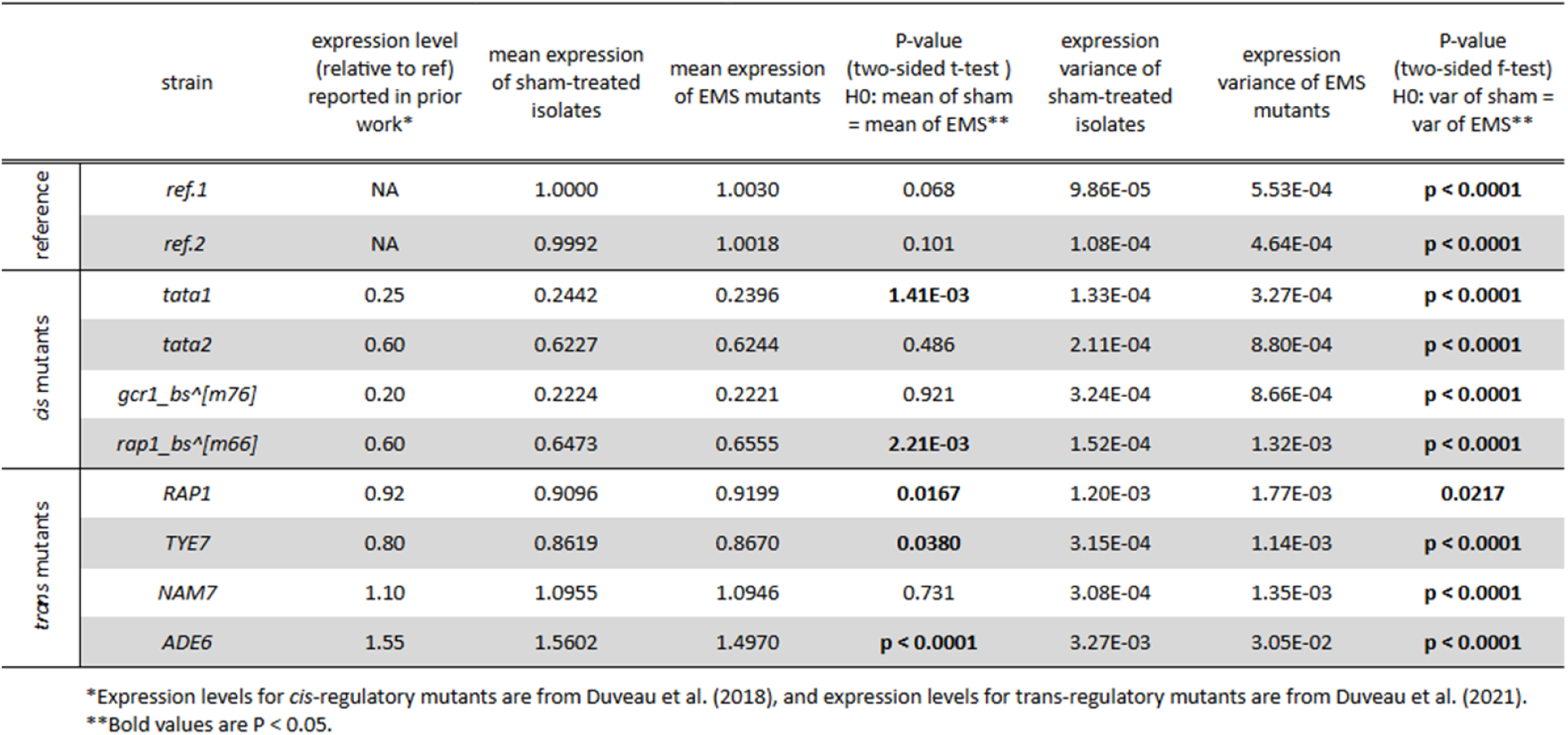
Effectsof new mutationson expression of the focal gene.

|  | strain | expression level<br>(relative to ref)<br>reported in prior<br>work* | mean expression<br>of sham-treated<br>isolates | mean expression<br>of EMS mutants | P-value<br>(two-sided t-test )<br>H0: mean of sham<br>= mean of EMS** | expression<br>variance of<br>sham-treated<br>isolates | expression<br>variance of EMS<br>mutants | P-value<br>(two-sided f-test)<br>H0: var of sham =<br>var of EMS** |
| --- | --- | --- | --- | --- | --- | --- | --- | --- |
| reference | *ref.1* | NA | 1.0000 | 1.0030 | 0.068 | 9.86E-05 | 5.53E-04 | **p < 0.0001** |
|  | *ref.2* | NA | 0.9992 | 1.0018 | 0.101 | 1.08E-04 | 4.64E-04 | **p < 0.0001** |
| *cis* mutants | *tata1* | 0.25 | 0.2442 | 0.2396 | **1.41E-03** | 1.33E-04 | 3.27E-04 | **p < 0.0001** |
|  | *tata2* | 0.60 | 0.6227 | 0.6244 | 0.486 | 2.11E-04 | 8.80E-04 | **p < 0.0001** |
|  | *gcr1_bs^[m76]* | 0.20 | 0.2224 | 0.2221 | 0.921 | 3.24E-04 | 8.66E-04 | **p < 0.0001** |
|  | *rap1_bs^[m66]* | 0.60 | 0.6473 | 0.6555 | **2.21E-03** | 1.52E-04 | 1.32E-03 | **p < 0.0001** |
| *trans* mutants | *RAP1* | 0.92 | 0.9096 | 0.9199 | **0.0167** | 1.20E-03 | 1.77E-03 | **0.0217** |
|  | *TYE7* | 0.80 | 0.8619 | 0.8670 | **0.0380** | 3.15E-04 | 1.14E-03 | **p < 0.0001** |
|  | *NAM7* | 1.10 | 1.0955 | 1.0946 | 0.731 | 3.08E-04 | 1.35E-03 | **p < 0.0001** |
|  | *ADE6* | 1.55 | 1.5602 | 1.4970 | **p < 0.0001** | 3.27E-03 | 3.05E-02 | **p < 0.0001** |

\*Expression levels for *cis*-regulatory mutants are from Duveau et al. (2018), and expression levels for *trans*-regulatory mutants are from Duveau et al. (2021).
\*\*Bold values are P < 0.05.

The distribution of expression levels for the 229-312 EMS-treated isolates derived from mutagenesis of each regulatory mutant or replicate of the reference strain (**Figure 2C**) was assumed to have expression noise similar to the corresponding sham population plus an increase in expression variance caused by the new EMS-induced mutations. Indeed, all eight regulatory mutant strains (and both replicates of the reference strain) showed a significantly larger variance in *P_TDH3_-YFP* expression among the EMS mutants than the sham-treated isolates (**Table 1**, two-sided *f*-test, H_0_: 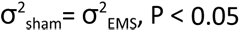). If the frequency and effects of new mutations increasing or decreasing expression of the focal gene were symmetric, the mean expression level of the EMS mutants should be similar to the mean expression level of the sham isolates. We found, however, that five of the eight regulatory mutant strains (*tata1*, *rap1_bs^m76^*, *RAP1*, *TYE7*, and *ADE6*) showed a statistically significant difference in mean expression level between the corresponding sham- and EMS-treated populations (**Table 1**, two-sided *t*-test, H_0_: μ_sham_=μ_EMS_, P < 0.05), suggesting that the effects of new mutations in these genetic backgrounds were asymmetric.

To infer strain-specific (or replicate-specific, for the reference strain) distributions of mutational effects from these data in a way that makes the distributions comparable among strains, we used the mean (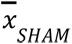) and standard deviation (σ*_SHAM_*) of the 99-157 sham-treated isolates from each of the eight mutant strains and two replicates of the reference strain to normalize the *P_TDH3_-YFP* expression level (*x_EMS_*) of each of the 229-312 EMS mutants isolated from the corresponding mutagenized population using the following formula (**Figure 2A**):

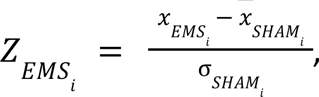

where *i* indicates the specific regulatory mutant strain or replicate of the reference strain. This calculation assumes the same expression noise for the corresponding EMS- and sham-treated populations and converts the expression level of each EMS mutant into a value similar to a Z-score, which is the term we will use for simplicity; the only difference is that Z-scores conventionally standardize differences from the mean value using the standard deviation from the same population. Our Z-scores describe changes in expression attributable to new mutations in units of standard deviation of the unmutagenized sham population for the same genotype. Each strain’s distribution of Z-scores is considered here to be an approximation of (and hereafter referred to as) that strain’s distribution of mutational effects (**Figure 2A**). The distributions of mutational effects observed for the eight regulatory mutant strains and the *ref.2* sample of the reference strain are shown in **Figure 2D**.

### Epistatic impacts of regulatory mutations on variance and skewness of distributions of mutational effects

Differences in the shape of the distribution of mutational effects reveal differences in the extent and range of phenotypic variation introduced by new mutations. Because each of the eight regulatory mutant strains examined in this study differed from the reference strain by a single regulatory mutation, we attribute any significant differences in the shape of the distribution of mutational effects between a regulatory mutant strain and the reference strain to epistatic interactions between the initial regulatory mutation carried by that strain and the new mutations introduced by EMS. To test for such epistatic effects, we compared the variance and skewness of the distribution of mutational effects for each regulatory mutant strain to the variance and skewness of the distribution of mutational effects for the reference strain.

The variance of a distribution of mutational effects is important because it measures mutational robustness, which is the sensitivity of a focal phenotype to the effects of new mutations (Félix and Barkoulas 2015). [This variance is related to, but distinct from, the mutational variance, *V_M’_*, often reported in quantitative genetics (see **Supplementary Note, Supplementary Table S3**).] Regulatory mutant strains showing a greater variance in their distribution of mutational effects than the reference strain are inferred to carry an initial regulatory mutation that interacts epistatically with new mutations in a way that makes them more likely to alter expression of the focal gene, decreasing mutational robustness. Because new mutations arising in such strains generate more variance in expression of the focal gene, such initial regulatory mutations can also be said to increase evolvability (Flatt 2005). The opposite is true for regulatory mutations in strains showing less expression variance in response to new mutations than the reference strain.

The skewness of a distribution of mutational effects is also important because the symmetry of a distribution of mutational effects determines whether new mutations are more likely to increase or decrease a trait’s value (e.g., a gene’s expression level). Traits with a right-skewed distribution of mutational effects are expected to increase the trait’s value in the absence of natural selection, whereas traits with a left-skewed distribution are expected to decrease the trait’s value. We used the *medcouple* statistic to measure skewness, which compares the scaled median difference between the left and right halves of a distribution and provides an estimate of skewness that is robust to outliers (Brys et al. 2004). Like all skewness metrics, *medcouple* ranges from -1 to +1 and is positive for right-skewed distributions and negative for left-skewed distributions.

To determine whether any observed differences in variance or skewness between a regulatory mutant strain and the reference strain are likely to have arisen by chance, we used a larger, previously published, dataset containing EMS- and sham-treated isolates for the same reference strain used in the current study to construct a null distribution for each parameter. This dataset (*ref.0*) included measures of *P_TDH3_-YFP* expression in 1213 EMS mutants and 146 sham-treated isolates (Metzger et al. 2016) (**Figure 3A, Supplementary Table S2**). These EMS mutants were collected following the same EMS mutagenesis and FACS sorting protocols described above and have a similar number of new mutations per mutant (**Supplementary Table S1**). To construct the null distribution for variance, we randomly sampled (with replacement) 300 of these 1,213 EMS mutants, used the mean and standard deviation of the *ref.0* sham isolates to convert the expression level of each of these EMS mutants into a Z-score, and calculated the variance of the resulting distribution of mutational effects (**Figure 3A**). We repeated this process 10,000 times, recording the variance of each random sample of 300 EMS mutants. These 10,000 measures of variance were used to define the null distribution for variance of the distribution of mutational effects for the reference strain. The same resampling process was employed to produce a null distribution for skewness (**Figure 3A**).

**Figure 3.**
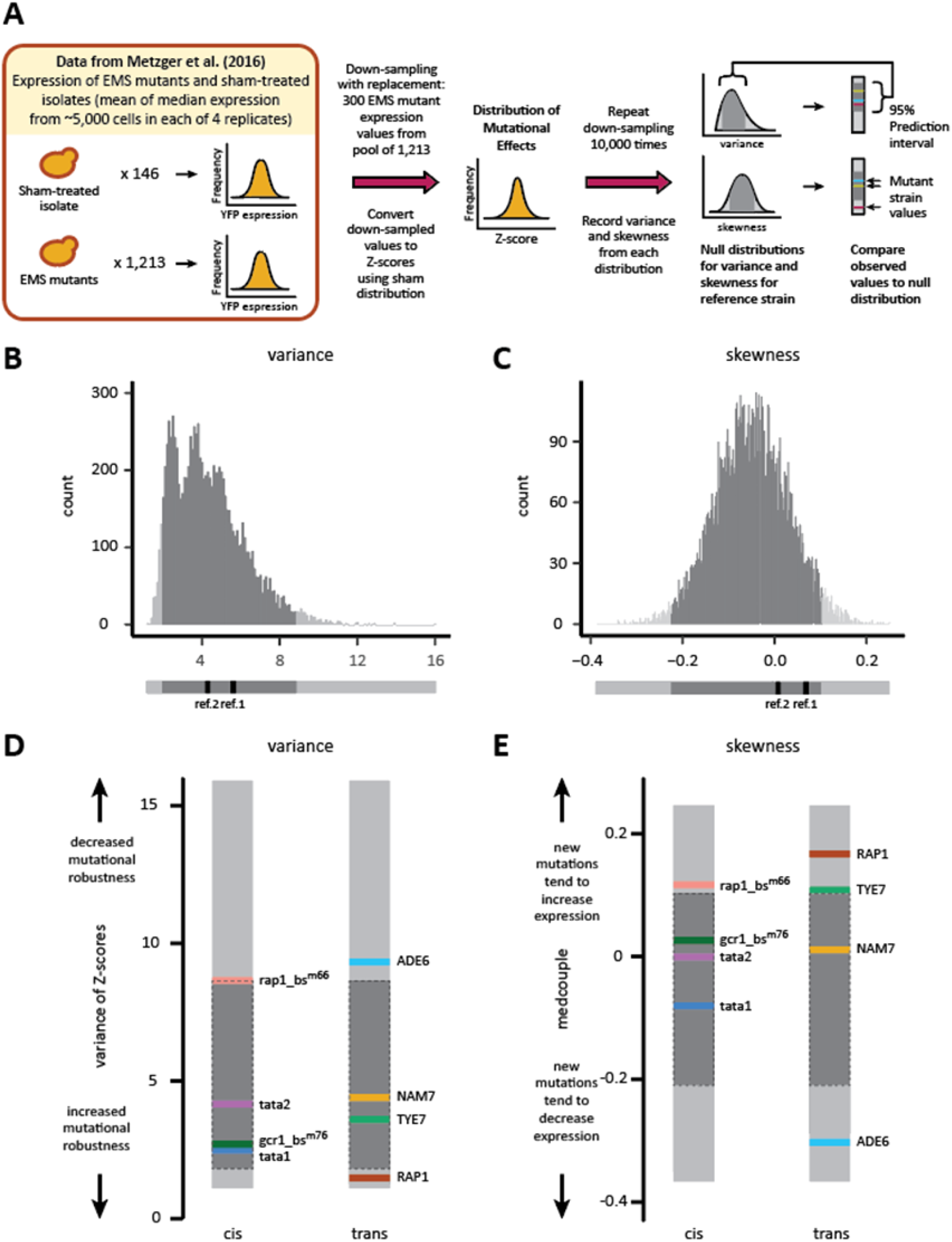
*cis*- and *trans*-regulatory mutations can alter the variance or skewness of the distribution of mutational effects for focal gene expression. (**A**) The schematic shows how we used expression data from 1213 EMS mutants and 146 sham-treated isolates reported in Metzger et al. (2016) that were derived from the same reference strain used in the current study and generated following the same protocols to construct null distributions for the variance and skewness of the distribution of mutational effects for the reference strain. Briefly, we sampled (with replacement) 300 EMS mutants from this set of 1213 mutants and used the mean and standard deviation of the 146 sham-treated isolates to convert the expression level of each of the 300 sampled EMS mutants into Z-scores. We then calculated the variance and skewness from the resulting distribution of mutational effects. We repeated this process 10,000 times, using the 10,000 measures of variance and skewness to form null distributions for the reference strain. The variance and skewness observed for the distributions of mutational effects defined by the 229-312 EMS mutants isolated from each of the eight regulatory mutant strains in the current study were then compared to these null distributions. Observed values that fell outside the 95% prediction interval for variance or skewness of the reference strain were interpreted as having come from an underlying distribution of mutational effects with a significant difference in variance or skewness, respectively, from the reference strain. (**B,C**) The null distributions obtained for the variance (**B**) and skewness (**C**) of the distribution of mutational effects from the reference strain are shown with the 95% prediction interval shaded dark gray. Below each histogram is an alternative line representation of the null distribution that shows the 95% prediction interval and the values of variance and skewness for *ref.1* and *ref.2*, the two replicate samples produced as part of the current study. (**D,E**) The observed values of variance (**D**) and skewness (**E**) for the distribution of mutational effects for each of the four *cis*- and four *trans*-regulatory mutant strains are shown on the line representation of the null distribution for both variance and skewness.

Differences in variance or skewness between one of the eight regulatory mutants and the reference strain were considered statistically significant if the observed value of variance or skewness for that regulatory mutant strain fell outside the 95% bootstrap prediction interval defined by the null distribution for the reference strain (**Figure 3A**). Both the variance and skewness observed for *ref.1* and *ref.2* fell within the 95% prediction interval (**Figure 3B and C**, **Table 2**), suggesting that the distributions of mutational effects described in this study are reproducible and robust to the specific new mutations sampled.

**Table 2.** Effects of cis- or trans-regulatory mutations on the variance and skewness of the distributions of mutational effects.

|  | strain | variance of Z-scores | Proportion of ref.0<br>bootstrap variance values<br>< mutant variance value | skewness of Z-scores<br>(medcouple) | Proportion of ref.0<br>bootstrap medcouple<br>values < mutant<br>medcouple value |
| --- | --- | --- | --- | --- | --- |
| reference | *ref.0* | 4.3559 | -- | -0.0569 | -- |
|  | *ref.1* | 5.6118 | 0.7726 | 0.0680 | 0.9338 |
|  | *ref.2 (shown in Fig. 2)* | 4.2813 | 0.5435 | 0.009009 | 0.7762 |
| cis mutants | *tata1* | 2.4576 | 0.1509 | -0.0804 | 0.3572 |
|  | *tata2* | 4.1631 | 0.5219 | -0.0006 | 0.7373 |
|  | *gcr1_bs^[m76]* | 2.6769 | 0.2001 | 0.0260 | 0.8299 |
|  | *rap1_bs^[m66]* | 8.7377 | 0.9741 | **0.1168** | **0.9821** |
| trans mutants | *RAP1* | **1.4783** | **0.0024** | **0.1670** | **0.9968** |
|  | *TYE7* | 3.5999 | 0.3885 | **0.1084** | **0.9767** |
|  | *NAM7* | 4.3953 | 0.5647 | 0.0110 | 0.7834 |
|  | *ADE6* | **9.3201** | **0.9846** | **-0.3050** | **0.0005** |

\*Bold values fall outside the 95% bootstrap prediction intervals

We found that two of the *trans-*regulatory mutant strains—(*RAP1* and *ADE6*)—showed a statistically significant difference in variance from the reference strain (**Figure 3D**, **Table 2**), indicating that epistatic interactions between new mutations and the initial regulatory mutation carried by each of these strains changed mutational robustness. The *RAP1* mutant strain showed significantly less variance resulting from the effects of new mutations than the reference strain, suggesting that the nonsynonymous mutation in the *RAP1* coding sequence increased mutational robustness. By contrast, the *ADE6* mutant strain showed increased variance attributable to the effects of new mutations, suggesting that the nonsynonymous mutation in the coding sequence of *ADE6* decreased mutational robustness.

The same two *trans-*regulatory mutant strains, plus one additional *trans*-regulatory mutant (*TYE7*) and one of the *cis-*regulatory mutants (*rap1_bs^m66^*) showed a statistically significant difference in skewness from the reference strain (**Figure 3E**, **Table 2**), suggesting a change in the direction and/or magnitude of mutational effects. The distribution of mutational effects was significantly more left-skewed (negatively skewed) for the *ADE6* mutant strain than the reference strain, meaning that the mean effect of new mutations in this strain decreased expression more than the median effect of new mutations in this strain. Indeed, mutations causing large decreases in expression were much more common in the distribution of mutational effects for the *ADE6* strain than the reference strain (**Figure 2D**). By contrast, the *TYE7*, *RAP1*, and *rap1_bs^m66^* strains were all significantly more right-skewed (positively skewed) than the reference strain (**Figure 3E**), consistent with mutations increasing expression having larger average effects than mutations decreasing expression (**Figure 2D**).

### Conclusions

We found that four of the eight regulatory mutations tested interacted epistatically with new mutations in a way that altered the shape of the distribution of mutational effects for expression of a focal gene. More specifically, we found that one of the four *cis*-regulatory mutations and three of the four *trans*-regulatory mutations tested altered the variance and/or skewness of the distributions of mutational effects inferred using point mutations introduced randomly throughout the genome by the chemical mutagen EMS. It is intriguing that we observed epistatic effects more often in the case of *trans-*regulatory mutations, as such a pattern might be expected given that mutations with *trans-*regulatory effects on expression of a focal gene tend to interact (directly or indirectly) with more genes in the genome than those in *cis* (Vande Zande and Wittkopp 2022). However, given the small number of mutants we examined here and the differences in effect sizes between the *cis-* and *trans-*regulatory mutants, the current data only allows us to speculate in this regard.

Regardless of whether this difference in epistatic impacts between *cis*- and *trans*-regulatory mutations emerges as a general trend after further investigation, our data demonstrate that as little as one nucleotide change in a regulatory network can be sufficient to alter the distribution of mutational effects for expression of a focal gene. This observation suggests that distributions of mutational effects for expression of the same gene might often differ among genotypes even *within* a species, indicating that caution is warranted when extrapolating the effects of new mutations from one genotype to the next. Indeed, a recent study showed that the effects of gene deletions on expression of other genes often differed between two strains of *S. cerevisiae* (Liu et al. 2024).

Perhaps even more intriguing is that all four regulatory mutations that caused a significant change in the skewness of the distribution of mutational effects for expression of our focal gene did so in a way that makes new mutations more likely to have effects that compensate for the initial regulatory mutation. For example, the *rap1_bs^m76^* mutation in the TDH3 promoter *decreased* its activity, but also altered the regulatory network in a way that caused new mutations to be more likely to *increase* its activity (**Figure 4A**). Nonsynonymous mutations in the *RAP1* and *TYE7* coding sequences also caused decreased expression driven by the *TDH3* promoter and biased the distribution of mutational effects in a way that makes new mutations more likely to increase expression (**Figure 4B,C**). Finally, the *ADE6 trans*-regulatory mutation increased expression driven by the *TDH3* promoter but also made new mutations more likely to decrease its expression (**Figure 4D**). This pattern is reminiscent of the diminishing returns epistasis often observed when studying the effects of new mutations on fitness and suggests that phenotype-correlated epistatic effects might be at play for traits other than fitness. It also suggests that there might be a selectively neutral mechanism favoring the production of compensatory mutations at the level of gene expression that could slow expression divergence and would be consistent with the surprisingly high frequency of compensatory mutations observed for activity of the *TDH3* reporter among naturally occurring strains of *S. cerevisiae* (Metzger and Wittkopp 2019).

**Figure 4.**
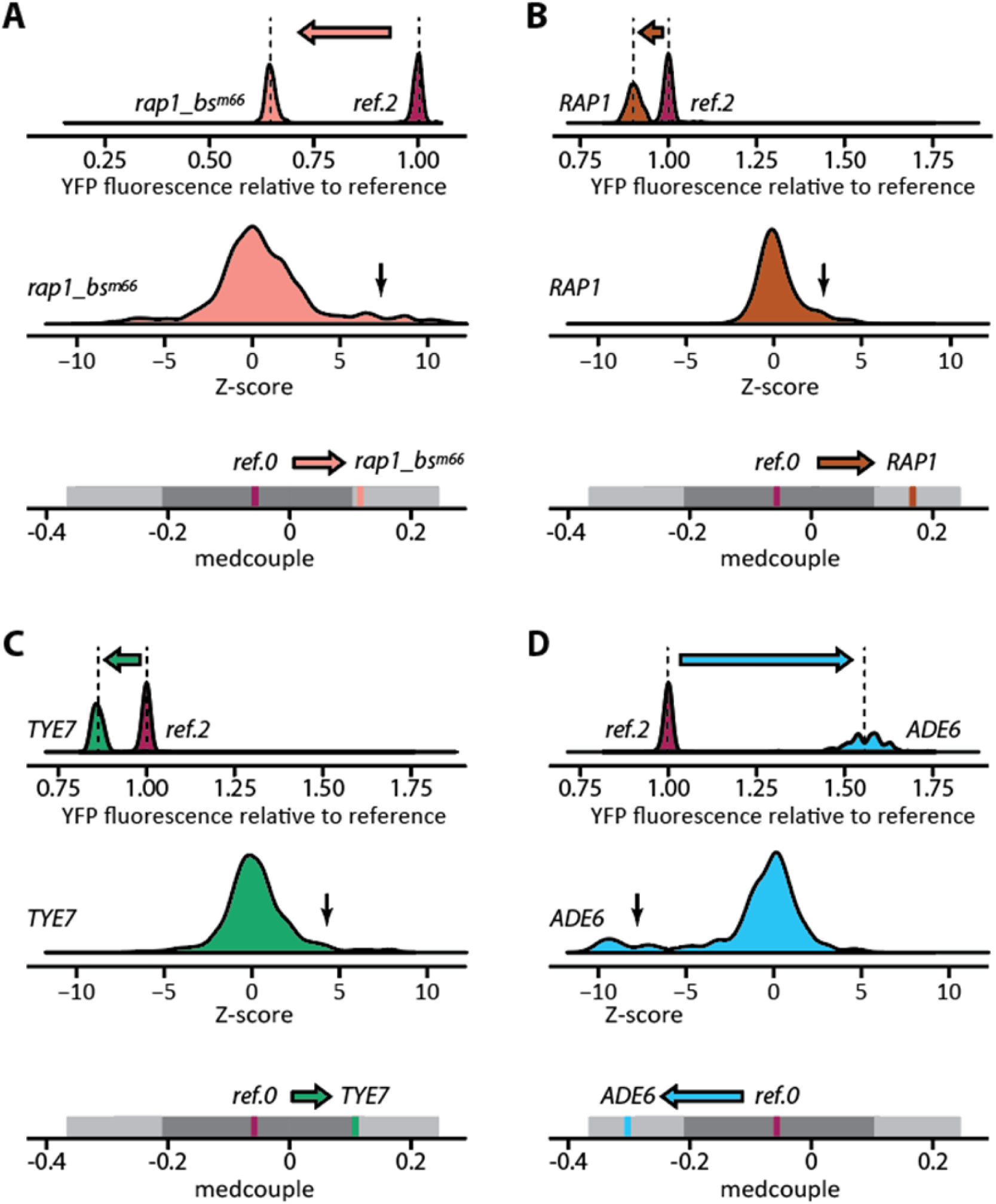
Epistatic interaction with regulatory mutations might reduce the probability of expression divergence. Data are shown for the *rap1_bs^m66^ cis*-regulatory mutant (**A**), the *RAP1 trans*-regulatory mutant (**B**), the *TYE7 trans*-regulatory mutant (**C**), the *ADE6 trans*-regulatory mutant (**D**), suggest that the single regulatory mutation distinguishing the regulatory mutant strain from each reference strain causes a change in skewness of the distribution of mutational effects for new point mutations that increases the chance that new mutations (at least partially) compensate for the effects of the initial regulatory mutation. In each panel, the top row shows distributions of expression levels for the *P_TDH3_-YFP* reporter gene for sham-treated isolates from the regulatory mutant strain and the *ref.2* sample of the reference strain. The arrow indicates the direction of change in expression caused by the single regulatory mutation introduced into the reference strain to make that regulatory mutant strain. The middle row in each panel shows the distribution of Z-scores for the EMS mutants obtained for the regulatory mutant strain, with a black arrow pointing out the heavy tails of the distribution. The impact of these heavy tails on skewness is reflected in the bottom row in each panel, which shows a line representation of the null distribution for skewness of the reference strain (*ref.0*), with the value observed for the regulatory mutant strain and the mean value observed for the *ref.0* strain indicated. The arrow in the bottom row of each panel shows the direction of change in skewness of the distribution of mutational effects caused by the initial regulatory mutation present in that regulatory mutant strain.

## Materials and Methods

### Yeast Strains

All strains used in this study were derived from strain YPW1139 (*Matα ura3d0*), which was constructed as described in Metzger et al. (2016). Relative to the standard BY laboratory strain, YPW1139 contains five genetic variants that decrease the frequency of petite mutants and increase the sporulation rate. It also contains a reporter gene inserted at the *HO* locus on chromosome IV. This reporter gene consists of the *TDH3* promoter allele from the BY laboratory strain driving expression of the coding sequence for a yellow fluorescent protein (YFP), followed by the *CYC1* terminator sequence and a KanMX4 drug resistance cassette.

The four *cis-*regulatory mutant strains (*tata1*, *tata2*, *gcr1_bs^m76^*, and *rap1_bs^m66^*) were generated by using site directed mutagenesis to introduce a single point mutation into the reference (BY) allele of the *TDH3* promoter of the reporter gene in YPW1139, as described in Metzger et al. (2015) and Duveau et al. (2018). These point mutations were located 505 bp (*rap1_bs^m66^*, strain YPW1897, G-505a), 482 bp (gcr1_bs^m76^, strain YPW1960, C-482t), 137 bp (*tata1*, TATA.81, strain YPW2605, T-137c, TATA**<u>C</u>**AAA) and 136 bp (*tata2*, TATA.99, strain YPW2622, A-136g, TATAT**<u>G</u>**AA) upstream of the *TDH3* start codon, and were reported to cause the *TDH3* promoter to have 60%, 20%, 25%, and 60%, respectively, of the activity of the unmutated reference *TDH3* promoter allele.

The non-synonymous mutations in *trans*-regulatory mutants *ADE6* (strain YPW2921, G3443a, changes codon 352 of 1359 from Gly to Asp), *TYE7* (strain YPW2728, C391t, changes codon 131 of 292 to a stop codon) and *NAM7* (strain YPW2800, G1304a, changes codon 435 of 972 from Gly to Asp) were originally isolated from the EMS mutagenesis screen described in Metzger et al. (2016), mapped using the methods described in Duveau et al. (2014) and Duveau et al. (2021), and reintroduced into the reference strain as described in Duveau et al. (2021). These mutations were reported in Duveau et al. (2021) to cause activity of the *TDH3* promoter that was 155%, 80%, and 110%, respectively, of the level of the unmutated reference promoter allele. The *trans-*regulatory mutant allele of *RAP1* used in this study was isolated from a deep mutational scanning experiment using error-prone PCR to create a library of mutant *RAP1* alleles with mutations in the coding sequence, as described in Duveau et al. (2021). The specific RAP1 mutant strain used here (strain YPW3287, which is listed as R357 in Duveau et al. (2021) supplementary file 16 and RAP1357 in Vande Zande et al. (2022) supplementary data S1) carries mutation G1918 in the RAP1 coding sequence that changes codon 640 out of 827 from D to N as well as two other synonymous mutations (T321c and A2415g). This RAP1 allele was reported to reduce *TDH3* promoter activity to 92% of that seen in the reference strain under the saturated growth conditions used here and in Duveau et al. (2021). Interestingly, when assayed during log-phase growth in Vande Zande et al. (2022), this RAP1 allele caused the *P_TDH3_-YFP* reporter gene to be expressed at 116% of the expression level seen for the unmutated reference strain, which was reported as log_2_ fold change = 0.21 in supplementary data S1 of Vande Zande et al. (2022).

### EMS mutagenesis

EMS mutagenesis was used to introduce new point mutations randomly throughout the genome, following the protocol described in Metzger et al. (2016). Briefly, before each mutagenesis experiment, yeast cells from each strain were revived from glycerol stocks stored at -80 °C on YPG agar medium (10g/L yeast extract, 20g/L peptone, 5% vol/vol glycerol and 20g/L agar) incubated for 48 hours at 30°C. The use of this medium minimizes the formation of spontaneous petite mutants because they are unable to grow on non-fermentable carbon sources such as glycerol. Cells from these solid plates were transferred into 10 ml YPD liquid medium (20g/L monosaccharides, 10g/L yeast extract and 20g/L peptone) and cultured for 10-11 cell cycles (∼24 hours) at 30 °C with shaking at 200 rpm. Before mutagenesis, cells were washed in 1 ml 1X PBS and 1 ml water twice and re-suspended in 1 ml of sodium phosphate (0.1M). 10μl EMS was then added to the suspension, resulting in an EMS concentration of 1%. After 45 minutes, EMS mutagenesis was quenched by adding 1 ml of 5% sodium thiosulfate. Cells were then washed twice with 1ml 5% sodium thiosulfate, followed by twice more with 1 ml water, in 1.7mL centrifuge tubes (Eppendorf). Next, cells were washed once in YPD and then resuspended in 1ml YPD. Finally, 0.125mL of this liquid culture was transferred to 3.875 mL of fresh YPD liquid medium. After 24 hours at 30 °C, 0.125 ml of each 4mL culture was diluted into 3.875 ml fresh YPD. These two cycles of dilution and growth allowed the cells to recover from the stress of the EMS treatment for approximately 10 cell cycles without overgrowth. Control, sham-treated, cells for each strain were treated with all wash and dilution steps described above, omitting only the addition of EMS.

The 1% concentration of EMS and 45 minute exposure time used in this experiment were the same conditions used in Metzger et al. (2016). These conditions were determined empirically in Gruber et al. (2012) to try to maximize the number of mutants recovered with statistically significant changes in *P_TDH3_-YFP* expression while minimizing the probability such changes in reporter gene expression were caused by more than one mutation. Using a canavanine resistance assay (described below), Metzger et al. (2016) estimated that these EMS mutagenesis conditions introduced an average of 32 (21–43, 95% confidence interval) new mutations per cell in the *ref.0* dataset (**Supplementary Table S1**). Subsequent genome sequencing, genetic mapping, and functional testing of 76 of the EMS mutants isolated in Metzger et al. (2016) showed that these EMS mutants actually carried an average of only 24 new point mutations per cell, suggesting that the canavanine resistance assay overestimated the mutation rate by 25% (Duveau et al. 2021). With few exceptions, a significant change in *P_TDH3_-YFP* expression observed in an EMS mutant was caused by only one of the approximately 24 mutations induced by EMS in that mutant (Duveau et al. 2021). Given the 12 Mb haploid genome size of *S. cerevisiae,* and assuming the spontaneous single nucleotide mutation rate of about 1.217 x 10^-10^ per base per generation (estimated based on the 873 point mutations observed in 145 diploid mutation accumulation lines over 2063 generations in Zhu et al. (2014), we expect approximately 0.00146 single nucleotide mutations to arise spontaneously per haploid genome each generation. The average of 28 point mutations we estimated were introduced into each haploid cell by the EMS treatment used for this study is thus similar to the number of point mutations we would expect to see introduced spontaneously after 19,178 generations.

### Canavaine resistance assay

To estimate the number of new mutations introduced by the EMS treatment, we used a canavanine resistance assay, following the protocol described in Gruber et al. (2012). Briefly, for each EMS-treated culture, 0.1 ml of a 10X dilution of cells were plated on agar medium (7.1g bacto-yeast nitrogen base, 20g/L dextrose, 2g/L amino acid mix without arginine and 20g/L agar) with 60 mg/ml L-canavanine sulfate and without arginine. In parallel, 0.1 ml of 2x10^-4^ dilutions of cells were also plated on the same agar medium but without L-canavanine. After incubating all plates at 30°C for 48 hours, we counted the number of colonies visible on each plate. Colonies that formed on the L-canavanine plates contained cells carrying a canavanine resistance mutation. The number of colonies on plates without L-canavanine were used to estimate the concentration of cells in that culture. We then used this information to infer the number of mutations causing canavanine resistance, which results from a finite set of known mutations in the *CAN1* gene (Lang and Murray 2008).

### Fluorescence Activated Cell Sorting (FACS) and recovery of EMS mutants and sham-treated isolates

After the 48-hour recovery period following EMS or sham treatment, fluorescence activated cell sorting (FACS) was performed on a BD FACS Aria II (University of Michigan Flow Cytometer Core). Before sorting, a 0.5ml YPD liquid culture (∼1x10^7^ cells) of each sample was diluted in 1.5ml 1X PBS. Each sample was then run on the FACS machine at a flow rate of ∼15000 cells/s with gates set manually in the FACSDiva software to exclude most non-yeast events and cell aggregates. For each EMS-treated population of cells, 384 events (cells or similarly sized debris) were randomly sorted (i.e., without regard to YFP fluorescence) onto YPD agar plates in a 384-well format. For each sham-treated population, 192 events were sorted in the same manner. Solid plates containing sorted events were then incubated for 48 hours at 30°C. Each colony that formed on a plate containing cells sorted from the EMS-treated population was considered a unique EMS mutant. Each colony that formed on a plate containing cells sorted from the sham-treated population was considered a unique sham-treated isolate. The number of EMS mutants and sham-treated isolates obtained from each sample is shown in **Supplementary Table S2**.

Cells from each of these colonies were moved from the 384-well format on the solid YPD agar plate to one of four 96-well plates containing 0.5 mL YPD liquid medium in each well. In parallel, the reference strain, YPW1139, which was the progenitor strain for all regulatory mutants analyzed in this study, was revived from a glycerol stock stored at -80 °C by growing in liquid YPD and then added to a fixed set of 24 empty wells of each of these 96-well plates as a control. Data from replicates of this control strain (YPW1139) were used to correct for potential plate position effects in our final analysis. Note that these control cells were not treated with EMS nor subjected to a single cell bottleneck. A negative control strain YPW978, which does not have a *P_TDH3_-YFP* reporter gene or any other source of YFP fluorescence, was prepared in the same manner and added to two random wells of each plate. Data from replicates of this negative control strain were used to account for auto-fluorescence in our final analysis. All 96-well plates were incubated at 30 °C while shaking at 500 rpm for 24 hours. The next day, 100μl of the YPD culture from each well was mixed with 23 μL of 80% glycerol to make glycerol stocks in the 96-well plates, and the plates were stored at -80 °C.

To determine which, if any, of the isolated EMS mutants or sham-treated isolates had acquired a petite mutation (which affects expression of *P_TDH3_-YFP*), we transferred the cells remaining after making glycerol stocks into 96-well plates containing 0.5 mL of YPG agar medium (in which petites cannot grow) and incubated the plates for 48 hours at 30°C while shaking at 500 rpm. Overall, ∼0.5%-2% of samples failed to grow, suggesting that they were petite mutants. These strains were excluded from further analysis.

### Measuring P_TDH3_-YFP expression in EMS mutants and sham-treated isolates

To determine the *P_TDH3_-YFP* expression level for each of the 229-312 EMS mutants and 99-157 sham-treated isolates from each of the eight regulatory mutants and two replicate populations of the reference strain (*ref.1* and *ref.2*), we used a 96-well pin tool (V & P Scientific) to transfer cells from each of the frozen 96-well glycerol stock plates onto solid YPG agar plates and incubated the plates for 48 hours at 30°C. Using a 96-well pin tool again, we then created four replicate 96-well plates from each of these original 96-well plates, with each well in each plate containing 0.5 ml of YPD liquid medium. These plates were incubated at 30°C for 22-24 hours while shaking at 500 rpm to reach saturated growth, and then 13-15 μl of each liquid culture was transferred to a new 96-well plate and mixed with 0.5 ml 1X PBS. YFP fluorescence and forward scatter (a proxy for cell size) were then measured for cells in each well of each plate using a BD Accuri C6 flow cytometer (488nm laser for fluorescence excitation and 533/30nm optical filter for signal acquisition) equipped with a HyperCyt autosampler (Intellicyt Corp).

These flow cytometry data were analyzed as described in Duveau et al. (2018) using scripts available on github (https://github.com/eden-mcqueen/epistasis-mutagenesis). Briefly, clustering functions in the R package *flowClust* (Lo et al. 2009) and *flowCore* (Hahne et al. 2009) were used to filter out all events that did not appear to be single-cells based on the estimated height and area of forward scatter. YFP fluorescence was then scaled by cell size, which was estimated by forward scatter. Next, the rescaled YFP fluorescence signal was transformed to better estimate YFP mRNA expression levels based on data directly comparing the two measures in Duveau et al. (2018). Samples with less than 2000 events remaining after filtering steps were excluded from further analysis. For the remaining samples, we calculated the median and standard deviation of the transformed YFP fluorescence for cells from each well. We then used the data from the 24 control wells on each plate containing the YPW1139 progenitor strain to correct the YFP fluorescence levels for any plate position effects on fluorescence. A single expression level for each EMS mutant and sham-treated isolate was then calculated as the mean of the medians observed for the four replicate samples. These YFP expression levels, as well as the median standard deviation for the four replicate cultures of each EMS mutant and each sham-treated isolate, are summarized in **Supplementary Table S4.**

Two-sided *t*-tests and *f*-tests, respectively, were used to compare the mean and variance of *P_TDH3_-YFP* expression between the corresponding sets of EMS mutants and sham-treated isolates (**Table 1**). These analyses were performed using base functions in R (R Core Team 2021).

### Comparing the variance and skewness of distributions of mutational effects among strains

To estimate the effects of new mutations on *P_TDH3_-YFP* expression for each of the eight regulatory mutant strains and both replicates of the reference strain (*ref.1* and *ref.2*) in a way that makes the distributions of mutational effects compatible among strains, we transformed the expression of each EMS mutant into a Z-score using the mean and variance of the corresponding sham-treated isolates with the formula 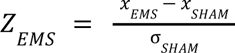, where *x_EMS_* is the expression level of an EMS mutant, and 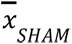 and *σ_SHAM_* are the mean and standard deviation, respectively, of expression of the corresponding sham-treated isolates. This transformation removes the different effects of the initial regulatory mutations on mean expression level and expression noise (measured as standard deviation) by expressing the effects of new mutations in units of standard deviation of the unmutagenized regulatory mutant or reference strain. A summary of these Z-scores for all EMS mutants and sham-treated isolates is included as **Supplementary Table S5**.

For each regulatory mutant strain and replicate of the reference strain, we calculated the variance of Z-scores using the base R function for variance (R Core Team 2021) and the skewness of Z-scores using the robust estimator *medcouple* as implemented in the *robustbase* R package (Maechler et al. 2023). To assess the significance of differences observed between the reference strain and each of the eight regulatory mutants, we generated null distributions for variance and skewness of the Z-score distribution for the reference strain by resampling from 1213 Z-score values derived from a large EMS mutant data set previously collected for this strain. Specifically, for each metric, we randomly sampled 300 Z-scores (with replacement) from this larger data set, calculated the value of the metric for the resultant Z-score distribution, and repeated this procedure 10,000 times. We then rank-ordered the 10,000 measures for each metric, and used the 250th and 9750th values in each set to define the lower and upper bounds, respectively, of a 95% bootstrap prediction interval for that metric in the reference strain. A regulatory mutant strain (or reference replicate) value for variance or skewness was considered significantly different from the reference strain if the value fell outside of this bootstrap prediction interval.

These 1213 EMS mutants, as well as the 146 sham-treated isolates that were used to convert the YFP expression level of each EMS mutant into a Z-score, were isolated and characterized in Metzger et al. (2016), using the same protocols used to isolate and characterize EMS mutants and sham-treated isolates in the current study. Each of these 1213 EMS mutants was derived from the 1340 events sorted randomly from the EMS-treated population by FACS in Metzger et al. (2016). The 146 sham-treated isolates were derived from the 160 events sorted from the sham-treated population in parallel. Z-scores for the 1213 EMS mutants in this *ref.0* dataset are included in **Supplementary Table S5**.

## Data Availability

A github repository is available for this manuscript at (https://github.com/eden-mcqueen/epistasis-mutagenesis). This repository includes R scripts (ANALYSIS.R, Clean.Data.R and Cleaning.Functions.2.R) and the Python script (create_template.py) used to process the raw .fcs flow cytometry data files analyzed in this study. Briefly, the Clean.Data.R script takes the .fcs files as input and uses the functions in “Cleaning.Functions.2.R” and the “create_template.py” script to produce an intermediate “Clean.Data2.txt” file. The ANALYSIS.R script takes this Clean.Data2.txt file as input and produces the “STRAIN.ESTIMATES.txt” data files, which are included in the .zip file. These STRAIN.ESTIMATES.#.txt files (where # ranges from 1 to 6) contain measures of YFP expression for each EMS mutant and each sham-treated isolate from each of the eight regulatory mutants as well as the *ref.1* and *ref.2* replicates of the reference strain. The SUMMARY.TRANS.txt data file is also included, which contains comparable data for the 1213 of EMS mutants and 146 sham-treated isolates from (Metzger et al. 2016). These data files include all the measurements used in the current manuscript, but also contain measurements for other types of samples collected in parallel for use in other studies. The “Analysis.1.9.R” script takes the STRAIN.ESTIMATES.x.txt files and the SUMMARY.TRANS.txt as input, selects the subset of data needed for the current study, adds more user readable names to the EMS mutants and sham-treated isolates, and performs the analyses presented in the manuscript. The raw flow cytometry data .fcs files generated for the *cis*-regulatory mutant strains is available at http://flowrepository.com (accession numbers: <pending>), and can be used to evaluate our data quality and methods. Unfortunately, due to a hard drive failure, the raw .fcs files for the *trans*-regulatory mutant strains and reference strain replicates are no longer available.

## Acknowledgements

We thank past and present members of the Wittkopp lab, including Fabien Duveau, Brian Metzger, David Yuan, Andrea Hodgins-Davis, Taslima Haque, Mohammad Siddiq, and Anna Redhuis for valuable discussions and input on this work. We also thank the University of Michigan Center for Chemical Genomics and the Flow Cytometry Core for access to flow cytometers. Funding for this work was provided by the National Institutes of Health (R35GM118073 to PJW) and the National Science Foundation (MCB 1929737 to PJW and DBI 2109787 to EWM). The content is solely the responsibility of the authors and does not necessarily represent the official views of the National Institutes of Health or the National Science Foundation.

