## Supplementary Information for "Epistatic impacts of *cis*- and *trans*-regulatory mutations on the distribution of mutational effects for gene expression in *Saccharomyces cerevisiae*"

**Table S1 - Estimated number of mutations per EMS mutant isolated**

**Table S2 - Estimates of mutation number based on canavanine resistance assay**

**Table S3 - Mutational variance (Vm) for point mutations**

**Table S4 - Summary of YFP expression level and expression noise for EMS mutants and sham-treated isolates (file included separately)**

**Table S5 - Summary of Z-scores for EMS mutants from regulatory mutants and reference strain (file included separately)**

**Supplementary Note - Comparing the variance of a distribution of mutational effects to mutational variance ($V_M$)**

**Supplementary Table S1: Estimated number of mutations per EMS mutant isolated**

| | Strain | # of mutations estimated using canavanine resistance | 95% CI for # of mutations |
|---|---|---|---|
| reference | *ref.0* | 32 | (21, 43) |
| | *ref.1* | 43 | (35, 49) |
| | *ref.2* | 38 | (33, 42) |
| *cis* mutants | *tata1* | 41 | (36, 46) |
| | *tata2* | 37 | (31, 43) |
| | *gcr1_bs^[m76]* | 39 | (31, 47) |
| | *rap1_bs^[m66]* | 40 | (33, 47) |
| *trans* mutants | *RAP1* | 38 | (30, 46) |
| | *TYE7* | 34 | (28, 40) |
| | *NAM7* | 36 | (29, 43) |
| | *ADE6* | 39 | (31, 47) |

**Supplementary Table S1** shows estimates for the number of mutations per cell for each genotype, calculated from the results of canavanine resistance assays, as described in the main text. The 95% confidence intervals (95% CI) for these values are also given.

**Supplementary Table S2. Number of EMS-treated and sham-treated genotypes isolated from each strain**

| | Strain | sham isolate count | EMS-treated isolate count |
|---|---|---|---|
| reference | *ref.0* | 146 | 1213 |
| | *ref.1* | 152 | 275 |
| | *ref.2* | 152 | 280 |
| *cis* mutants | *tata1* | 157 | 290 |
| | *tata2* | 119 | 229 |
| | *gcr1_bs^[m76]* | 154 | 286 |
| | *rap1_bs^[m66]* | 119 | 229 |
| *trans* mutants | *RAP1* | 108 | 246 |
| | *TYE7* | 141 | 304 |
| | *NAM7* | 152 | 280 |
| | *ADE6* | 99 | 312 |

**Supplementary Table S2** lists the number of single colony isolates that were collected from each of the mutagenized and sham populations for each genotype and carried forward into subsequent analysis.

**Supplementary Table S3. Mutational variance (Vm) for mutant genotypes and reference strains**

| | Strain | expression variance in sham distribution (Ve) from Table 1 | expression variance in EMS distribution (Vp) from Table 1 | Mutational Variance (Vm) (assuming 1 generation) | Per Generation Mutational Variance (Vm) (assuming 19,178 generations of point mutations) | Mutational Heritability (Vm/Ve) (assuming 1 generation) | Mutational Heritability + 1 (compare to variance of Z-scores in Table 2) |
|---|---|---|---|---|---|---|---|
| reference | *ref.1* | 9.86E-05 | 5.53E-04 | 4.55E-04 | 2.37E-08 | 4.6118 | 5.6118 |
| | *ref.2* | 1.08E-04 | 4.64E-04 | 3.55E-04 | 1.85E-08 | 3.2813 | 4.2813 |
| *cis* mutants | *tata1* | 1.33E-04 | 3.27E-04 | 1.94E-04 | 1.01E-08 | 1.4576 | 2.4576 |
| | *tata2* | 2.11E-04 | 8.80E-04 | 6.68E-04 | 3.48E-08 | 3.1631 | 4.1631 |
| | *gcr1_bs^[m76]* | 3.24E-04 | 8.66E-04 | 5.43E-04 | 2.83E-08 | 1.6769 | 2.6769 |
| | *rap1_bs^[m66]* | 1.52E-04 | 1.32E-03 | 1.17E-03 | 6.12E-08 | 7.7377 | 8.7377 |
| *trans* mutants | *RAP1* | 1.20E-03 | 1.77E-03 | 5.74E-04 | 2.99E-08 | 0.4783 | 1.4783 |
| | *TYE7* | 3.15E-04 | 1.14E-03 | 8.20E-04 | 4.28E-08 | 2.5999 | 3.5999 |
| | *NAM7* | 3.08E-04 | 1.35E-03 | 1.05E-03 | 5.45E-08 | 3.3953 | 4.3953 |
| | *ADE6* | 3.27E-03 | 3.05E-02 | 2.72E-02 | 1.42E-06 | 8.3201 | 9.3201 |

**Supplementary Table S3** shows the values associated with calculation of mutational variance (*Vm*) for mutant and reference strains. As described in the **Supplementary Note** below, this value is distinct from the variance of the distribution of mutational effects referenced elsewhere in the manuscript. *Ve* represents the environmental variance, *Vp* represents the phenotypic variance, and *Vm* represents the mutational variance.

**Supplementary Table S4 - Summary of YFP expression level and expression noise for EMS mutants and sham-treated isolates**

**Supplementary Table S4** is included as a separate file and summarizes the YFP expression level (mean.medians) and expression noise measured as the standard deviation (median.sd) only for the EMS mutants and sham-treated isolates from the eight regulatory mutants and the three replicates (*ref.1*, *ref.2*, and *ref.0*) of the reference strain used in this current study. Note that the $gcr1\_bs^{m76}$ and $rap1\_bs^{m66}$ regulatory mutant strains are abbreviated as m76 and m66, respectively, in the table. The values of expression level (mean.medians) and expression noise (median.sd), respectively, come from the "YFP.MEDIAN.R2WT.MEAN" and "YFP.MEDIAN.R2WT.SD" measures in the STRAIN.ESTIMATES.x.txt files and the "YFP.MEDIAN.RELATIVE.MEAN" and the "YFP.MEDIAN.RELATIVE.SD" measures in the SUMMARY.TRANS.txt file.

**Supplementary Table S5 - Summary of Z-scores for EMS mutants from regulatory mutants and reference strain**

**Supplementary Table S5 is** included as a separate file and **s**ummarizes the Z-scores calculated from the data included in **Supplementary Table S4** for all the EMS mutants examined, including from the *ref.0* dataset.

**Supplementary Note: Comparing the variance of a distribution of mutational effects to mutational variance ($V_M$)**

In this study, we use the variance of the distribution of mutational effects as a measure of mutational robustness. This measure of variance is not the same as the mutational variance (VM) often used in quantitative genetic studies of polygenic traits, which describes the increase in phenotypic variance caused by the introduction of new mutations each generation (Lynch 1988). However, with a few assumptions, our data can be used to estimate VM for the reference strain and each of the eight regulatory mutant strains.

If we consider our EMS mutagenesis experiment a single generation, the phenotypic (gene expression) variance (VPEMS) observed for the 229-312 EMS-treated isolates from each strain is equal to the starting genetic variance (VG) present prior to mutagenesis plus the environmental variance (VE) and the mutational variance (VM) resulting from the introduction of new mutations (i.e., VPEMS = VG+VE+VM). Because the EMS mutagenesis was performed on a clonal starting population, we assume that VG = 0 and the equation for phenotypic variance in a population of EMS-treated isolates reduces to VPEMS =VE+VM.

The 99 to 157 sham-treated isolates from each strain were also derived from the same clonal starting population as the corresponding EMS-treated isolates, so VG = 0 in this case as well. Moreover, if we assume no new mutations arose in the sham-treated isolates, which prior work suggests is reasonable (Gruber et al. 2012; Metzger et al. 2016), VM= 0 for this population. Consequently, the phenotypic (gene expression) variance observed among the sham-treated isolates from the same strain reduces to VPSHAM =VE. The variance in gene expression observed among a set of sham-isolates is therefore taken as an empirical measure of VE for that strain under our experimental conditions. Assuming that VE is the same for the corresponding EMS- and sham-treated isolates, we can calculate the mutational variance for that strain as VM = VPEMS-VPSHAM because VM= (VM+VE)-VE.

But measures of VM are typically defined per generation, so to make this estimate more comparable to other studies, we must also scale this value by the number of generations it would take a haploid *S. cerevisiae* cell to acquire the ~28 spontaneous point mutations present in each of our EMS mutants. In a study of *S. cerevisiae* mutation accumulation lines, Zhu et al (Zhu et al. 2014) observed 873 point mutations among 145 diploid MA lines propagated asexually for 2062 generations, suggesting that a haploid S. cerevisiae genome should acquire an average of 873/145/2/2062 = 0.00146 point mutations per haploid genome per generation. Each of our haploid EMS-treated isolates is estimated to carry ~28 point mutations on average, suggesting that it carries a similar number of point mutations to those that would be spontaneously accumulated over (28/0.00146) = 19,178 generations. The values of VM estimated per generation for the eight regulatory mutant strains and two replicate samples from the reference strain analyzed in this experiment are shown in **Supplementary Table S3**. We note that these values are lower than estimates of VM for gene expression reported previously in *S. cerevisiae* based on genome-wide measures of gene expression and other mutation accumulation lines that included the effects of point mutations as well as structural variants and aneuploidies (Landry et al. 2007; Zhu et al. 2014).

Interestingly, although VM and the variance of our Z-score-based distributions of mutational effects are not the same, they are related through mutational heritability if our experiment is considered a single generation. Mutational heritability is a dimensionless metric that allows the phenotypic effects of new mutations to be compared among different traits and species (Houle et al. 1996); it is calculated by scaling VM by VE . The variance observed among the Z-scores defining our distributions of mutational effects is mathematically equivalent to the mutational heritability (VM/ VE) + 1 when the experiment is considered as a single generation (**Supplementary Table S3**).
